# Development of genomic resources for four potential environmental bioindicator species: *Isoperla grammatica*, *Amphinemura sulcicollis*, *Oniscus asellus and Baetis rhodani*

**DOI:** 10.1101/046227

**Authors:** Hannah C Macdonald, Luis Cunha, Michael W Bruford

## Abstract

A low-coverage genome was generated for each of four environmental key-species of macroinvertebrate taxa for the primary purpose of microsatellite marker development. *De novo* assemblies and microsatellite markers were designed for the freshwater species *Isoperla grammatica, Amphinemura sulcicollis,* and *Baetis rhodani* but have not been completed for the common shiny woodlice *Oniscus asellus*. Here, the data is made available, and the methods and pipeline are described which led to the creation of this resource. As widespread and functionally important organisms, which are often neglected in favour of studies on vertebrates, this data will be a useful resource for further research.

## Introduction

Macroinvertebrates are widespread, often dominant and functionally important members of their environment that, coupled with their relative ease of sampling make them ideally suited for use as indicator species for biomonitoring and conservation assessment (Pfrender et al. 2010; Buss et al. 2015; Cardoni et al. 2015), and are recognised as such by the Water Framework Directive (2000/60/CE) (European Commission 2000). However, their use in genetic approaches is still limited, often being neglected from studies because of the lack of data (Cardoso *et al*. 2011).

Four macroinvertebrate species were sequenced for the primary purpose of developing microsatellite markers for use in population genetics; these include three freshwater invertebrates (*Amphinemura sulcicnllis, Isoperla grammatica*, and *Baetis rhodani*) and one terrestrial soil invertebrate, the common shiny woodlice, *Oniscus asellus*. They all represent dominant, widespread species and therefore can be used as biomonitoring tools that will be effective at large spatial scales, as policy demands (Statzner and Bêche 2010). Microsatellite markers within this group are scarce, for example, within the large and diverse groups of Plecoptera and Ephemeroptera, there are only five species with between 3-13 microsatellites each, therefore this data will be a valuable and considerable resource for future research. This data could be used for further study into these invertebrates, such as describing their mitochondrial genome (as in Stewart and Beckenbach (2006)), or studying their genome content; evolutionary analyses (e.g. divergent rates), and further investigation of their genetic features (as in Li *et al*. (2010).

## Data ccess

Raw data is stored in NCBI’s Sequence Read Archive (SRA): NGS data for four invertebrates: Amphinemura sulcicollis, Isoperla grammatica, Baetis rhodani and Oniscus asellus (STUDY: PRJNA315680 (SRP072016)).

1. NGS sequence data (raw data sent from sequencing centres):

- *Amphinemura sulcicollis:* SAMPLE: Amphi_NGS (SRS1349204) EXPERIMENT: Amphi_NGS (SRX1642982) RUN: Amphi_NGS (SRR3262386)

WTCHG_93433_274_1.fastq.gz
WTCHG_93433_274_2. fastq.gz
WTCHG_93434_274_1.fastq.gz
WTCHG_93434_274_2. fastq.gz
- *Isoperla gramma tica:* SAMPLE: Iso_NGS (SRS1351356) EXPERIMENT: Iso_NGS (SRX1648180) RUN: Iso_NGS (SRR3262388)

WTCHG_93433_273_1.fastq.gz
WTCHG_93433_273_2.fastq.gz
WTCHG_93434_273_1.fastq.gz
WTCHG_93434_273_2.fastq.gz
- *Baetis rhodani:* SAMPLE: Baetis_NGS (SRS1351357) EXPERIMENT: Baetis_NGS (SRX1648181) RUN: Baetis_rhodani_NGS (SRR3262630)

Beatis_L3_1.fq.gz. 1.gz
Beatis_L3_1.fq.gz.2.gz
Beatis_L3_2.fq.gz. 1.gz
Beatis_L3_2.fq.gz.2.gz
- *Oniscus asellus:* SAMPLE: Woodlice_NGS (SRS1351401) EXPERIMENT: Oniscus_NGS (SRX1648318) RUN: Oniscus_NGS (SRR3263253)

WTCHG_93433_275_1.fastq
WTCHG_93433_275_2.fastq
WTCHG_93434_275_1.fastq
WTCHG_93434_275_2.fastq
2. Each freshwater species has a CONTIG file (after *de novo* assembly) deposited at DDBJ/ENA/GenBank under the accession’s listed below, any contigs under 200bp were removed.

- *Amphinemura sulcicollis:* SUBID: SUB1394726 BioSample: SAMN04568201 Accession: LVVV00000000 Organism: Amphinemura sulcicollis Dwyryd File name: amphi_kmer61.contig
- *Isoperla grammatica:* SUBID: SUB1397890 BioSample: SAMN04568202 Accession: LVVW00000000 Organism: Isoperla grammatica Teifi File name: iso_kmer61.contig
- *Baetis rhodani:* SUBID: SUB1398024 BioSample: SAMN04568203 Accession: LVVX00000000 Organism: Baetis rhodani Ty wi File name: B_61.contig
3. Other data stored in Genbank:

- 132 Mitochondrial cytochrome c oxidase I (mtCOI) sequences for all four species (using barcoding primers from Folmer *et al*. (1994)) are available on genbank: Accession numbers KU955863-KU955994 *(Amphinemura sulcicollis* 31 sequences; *Isoperla grammatica* 29 sequences; *Baetis rhodani* 65 sequences; *Oniscus asellus* 6 sequences).
- o 51 Microsatellite markers for *Isoperla grammatica*, *Amphinemura sulcicollis* and *Baetis rhodani* available on Genbank: between KR068997-KR069048 (Iso_1-18, Amp_1-21, B_1-13, respectively) and described fully in Macdonald *et al*. (2016) *in review*. Subsequent microsatellite marker development has not been completed for *Oniscus asellus*.

## Meta Information

Data for the four draft genomes sequenced (Table 1) was generated at two different sequencing centres, which were compared for their cost effectiveness and yields. Libraries 1-3 (*A. sulcicollis*, *I. grammatica and O. asellus*) were sent to Oxford MRC Sequencing, multiplexed with five other samples (eight libraries) as part of collaboration at Cardiff University. Whereas *B. rhodani* was sent at a later date to Beijing Genomic Institute (BGI), along with two other samples, these three samples were labelled by BGI and multiplexed in one lane (see Table 1 for full details). The main goals of the experiment were to develop enough genomic resources for each target species in order to retrieve enough high quality microsatellite markers.

**Table 1.** Details of the development of four separate libraries of macroinvertebrate using next generation sequencing.

|  | Library 1 | Library 2 | Library 3 | Library 4 |
| --- | --- | --- | --- | --- |
| Species | *Amphinemura sulcicollis* (Stephens, 1836) | *Isoperla grammatica* (Poda, 1761) | *Oniscus asellus* (Linnaeus 1758) | *Baetis rhodani* (Pictet, 1845) |
| Genus | Amphinemura | Isoperla | Oniscus | Baetis |
| Order | Plecoptera |  | Isopoda | Ephemeroptera |
| Class | Insecta |  | Malacostraca | Insecta |
| Meta Information |  |  |  |  |
| Sequencing centre | The Oxford Genomics Centre (WTCHG) / High-Throughput Genomics (Oxford, UK) |  |  | BGI (Shenzhen, China) |
| Platform | Illumina |  |  |  |
| Model | HiSeq 2500, Rapid run |  |  | HiSeq 2000 (PE91) |
| Analysis type | DNA |  |  |  |
| Run date | 10.12.2013 |  |  | 13.10.2014 |
| Library |  |  |  |  |
| Strategy | Whole-genome shotgun sequencing of genomic DNA |  |  |  |
| Shared lane | One lane (by itself) | One lane (by itself) | One lane (by itself) | One lane (shared with two other samples) |
| Sample type (mtDNA seq name available in genbank) | One individual (95A1) | One individual (96I1) | Two individuals mixed (71W3 and 70W1) | One individual (102B3) |
| Sex | Unknown |  |  |  |
| Source of material (Taxon) | Tissue |  |  |  |
| Sample Location# | 275070E 342810N | 268100E 254500N | 298106E 231045N<br>309487E 230414N | 273879E 246723N |
|  | Catchment: Dwyryd Upland Wales, UK | Catchment: Teifi Upland Wales, UK | Brecknock Wildlife trust reserve, Upland Wales, UK | Catchment: Tywi Upland Wales, UK |
| Insert length | 450 | 450 | 450 | 200 |
| Max read length | 300 | 300 | 300 | 200 |
| Results |  |  |  |  |
| Total reads (before QC) | 79,196,610 | 71,727,142 | 81,811,132 | 123,076,504 |
| Total reads (after QC) | 71,796,770 | 64,628,924 | 74,813,127 | 122,669,217 |
| % removed | 10.3 | 11.0 | 9.4 | 0.3 |

bioRxiv: Genomic Resource note
|  | **Library 1** | **Library 2** | **Library 3** | **Library 4** |
| --- | --- | --- | --- | --- |
| Species | *Amphinemura sulcicollis* (Stephens, 1836) | *Isoperla grammatica* (Poda, 1761) | *Oniscus asellus* (Linnaeus 1758) | *Baetis rhodani* (Pictet, 1845) |
| Mode of assembly | *De novo* assembly | *De novo* assembly | Flash | *De novo* assembly |
| Best kmer size | 61 | 61 | / | 61 |
| N50 | 1,543 | 568 | / | 850 |
| Total No. of contigs | 91,245 | 356,623 | / | 144,347 |
| Total scaffold length | 182,044,631 | 304,248,447 | / | 162,525,650 |
| Longest scaffold | 148,977 | 16,928 | / | 47,828 |

## Library

Multiple samples of each species were collected from sites around upland Wales, UK (Table 1) and stored in absolute ethanol. Genomic DNA was extracted from whole individuals using a High Pure PCR Template Preparation Kit for blood and tissue following the manufacturer’s instructions (Roche Diagnostics GmbH Mannheim, Germany). All samples were treated with RNase after DNA extraction. Individual samples were identified using Sanger sequencing, with standard barcoding primers from Folmer *et al*. (1994) and by comparing the sequences with data in Genbank. To assure sample quality, quantification was assessed using a Qubit and visualised on a gel (Figure 1). Nanodrop was used to assess contamination, where the 260/280 ratio were found to be between 1.8 and 2 and that the 260/230 ratio was between 2-2.2 across all analysed samples. The highest quantity and best quality samples were chosen; all species yielded DNA quantities required (which was 1-5μg of DNA normalized to a concentration of 50ng/μl) apart from *A. sulcicollis*, for which a vacuum concentrator had to be used. Samples showed high DNA integrity with no observed smearing on the electrophoresis gel (Figure 1).

**Figure 1.**
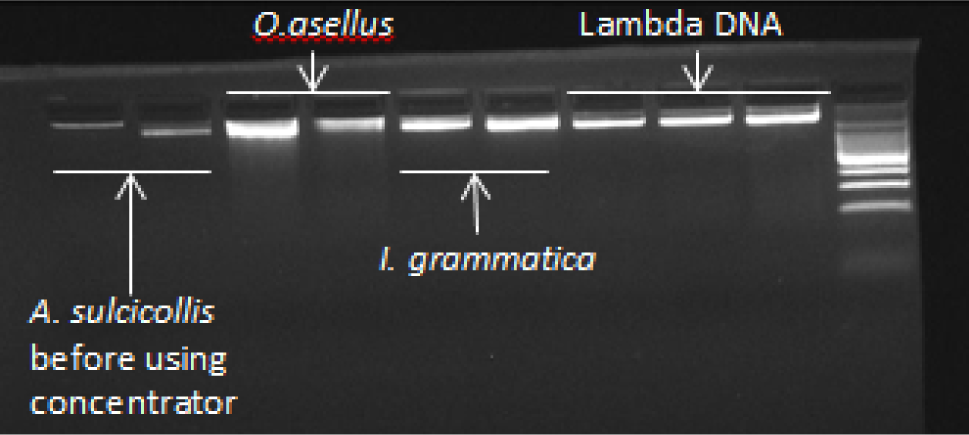
Photograph of a 3% ethidium-bromide stained agarose electrophoresis gel under UV light, showing genomic DNA of two individuals each species of the three species *Isoperla grammatica, Amphinemura sulcicollis* and *Oniscus asellus* that were sequenced first, compared to three concentrations of lambda DNA (left to right: 16.5ng/μL, 34 ng/μL and 67 ng/μL).

The samples of *A. sulcicollis, I. grammatica*, and *B. rhodani* were all made up of only one individual; however the *O. asellus* sample is made up of two individuals pooled. This was because allozyme loci have been used to show that two genetically distinct sub populations of *O. asellus* exist (*O. asellus* and *O. occidentails*) (Bilton *et al*. 1999) within *O. asellus*, it was thought that mixing two individuals would give the highest chance of success at developing microsatellites for the largest amount of samples. However, this meant that a *de novo* assembly could not be performed on this species.

Genomic DNA for all four samples were sent to their respective sequencing centres for library preparation (DNA was sheared, Illumina adapters were ligated, libraries were controlled for quality, normalized, pooled) and sequencing on HiSeq run (Table 1).

## Processing

For each library NGS created four raw Illumina read files (two libraries, each with two pairs), which was transferred to Linux, unzipped, and the two libraries were concatenated, leaving two files of two pairs (renamed from the raw file names in section Data Access, to Amp_1.fastq & Amp_2.fastq, Iso_1.fastq & Iso_2.fastq, Woo.fastq & Woo_2.fastq, and B_1.fastq & B_2.fastq for *I. grammatica, A. sulcicollis, O. asellus and B. rhodani*, respectively, Table 1). Quality control was preformed using Trimmomatic v0.32 (Lohse M *et al*. 2012) and Musket v1.1 Musket (Yongchao Liu *et al*. 2013). Trimmomatic was used to cut adapters and other illumina-specific sequences from the reads. It was also used to remove reads of low quality and short length. In this case the threshold for quality window was set at 18 and the minimum length was 35bp (using phred33). Musket is multistage k-mer based corrector for Illumina short read data and was used to identify and remove any common Illumina errors for a higher quality *de novo* genome assembly.

For all aquatic (single sample) species SOAP *de novo* 2 (Luo *et al*. 2012) was then used to build *de novo* assemblies using short-reads. This was done in order to provide longer reads for microsatellite marker mining. Several *de novo* assemblies were run per species in order to test different Kmer values and best assembly metrics. Draft assemblies were chosen according to maximum contig and highest N50 value.

A *de novo* assembly was not attempted for the pooled sample of *O. asellus* due to the risk of chimeras, which is much higher for assemblies of mixed samples. Instead FLASH v 1.2.9 (Fast Length Adjustment of SHort reads) was used to merge paired ends creating reads of 300 bp (http://ccb.jhu.edu/software/FLASH/MANUAL[Date accessed: 02.03.16]).

*PrimerPipeline* (http://www.scrufster.com/primerpipeline/ [Date accessed: 02.03.16]) was then used to identify repeat regions within the data files and design forward and reverse primers for each microsatellite. It is a windows program incorporating MISA (MIcroSAtellite identification tool, http://pgrc.ipk-gatersleben.de/misa/ [Date accessed: 02.03.16]) and Primer3 v.2.3.6 (Untergasser *et al*. 2012).

The full pipeline (including all scripts and annotations) is described in ‘Appendix 1 Script_NGS’ at the end of this manuscript.

## Results

The NGS for all four species was very successful as the total number of reads (raw data) were very high (ranging from 71,727,142 to 123,076,504 reads), and they were of relatively high quality because the quality control sections of the pipeline did not remove too much (ranging from 0.3% and 11% of the total reads, see Table 1). *B. rhodani* data from BGI appears to be the most successful as it had the highest total reads and the lowest percentage of reads removed by quality control.

The *de novo* assemblies varied according to which kmer size was used (see Table 2 for an example of how the kmer size affected the N50 in *A. sulcicollis*). For all three species that assemblies were performed for, kmer 61 was chosen as it produced the best assemblies. *A. sulcicollis* had the best N50 at 1,543, whereas *I. grammatica* had 568, meaning that on average the *de novo* assembly for *A. sulcicollis* produced larger contigs, therefore the best assembly.

**Table 2.** Shows information on all *de novo* assemblies preformed with *Amphinemura sulcicollis* data.

|  | Kmer_55 | Kmer_61 | Kmer_71 | Kmer_81 | Kmer_85 | Kmer_91 | Kmer_101 |
| --- | --- | --- | --- | --- | --- | --- | --- |
| Scaffold number | 88,343 | 91,245 | 89,130 | 87,731 | 88,028 | 87,347 | 92,636 |
| In-scaffold contig number | 834,812 | 832,323 | 832,173 | 842,707 | 850,059 | 864,994 | 1,170,180 |
| Total scaffold length | 184,467,063 | 182,044,631 | 180,880,141 | 183,600,553 | 181,179,087 | 179,755,162 | 172,156,150 |
| Average scaffold length | 2,088 | 1,995 | 2,029 | 2,092 | 2,058 | 2,057 | 1,858 |
| Filled gap number | 177,030 | 183,386 | 171,646 | 170,306 | 161,405 | 153,310 | 173,854 |
| Longest scaffold | 149,091 | 148,977 | 140,148 | 149,117 | 147,505 | 147,516 | 113,833 |
| Scaffold and singleton number | 606,125 | 580,009 | 605,194 | 626,038 | 645,302 | 671,404 | 950,155 |
| Scaffold and singleton length | 297,424,877 | 287,874,638 | 295,491,963 | 300,591,981 | 302,518,974 | 303,469,211 | 305,381,813 |
| Average length | 490 | 496 | 488 | 480 | 468 | 451 | 321 |
| N50 | 1,550 | 1,543 | 1,482 | 1,525 | 1,466 | 1,448 | 1,141 |
| N90 | 195 | 193 | 192 | 196 | 198 | 203 | 102 |
| Weak points | 0 | 0 | 0 | 0 | 0 | 0 | 0 |

## Acknowledgements

The research was funded by the NERC DURESS project (Diversity in Upland Rivers for Ecosystem Service Sustainability, NE/J01481/1) within the Biodiversity and Ecosystem Service Sustainability (BESS) Thematic Programme. HCM was funded by the Cardiff University President’s Research Scholarships. Many thanks to Steve Ormerod and Hefin Jones from Cardiff University, for support and guidance.

## Appendix 1: NGS script

Key: (Commands in red, Annotations and Instructions in Black)

### 1.0. Check and Unzip

#To check data files are unaffected from download/upload use md5sum to check the unique identity of the file (compare the md5 number, it has to be exactly the same format)

**md5sum filel > filel_md5.txt**

#And check the first line of the sequence

**head filename**

#E.g. md5sum formats for *B. rhodani* raw data files:

#aa278e7dd0de7af2e12aaf0d4ba9fc97 Beatis_L3_1.fq.gz.cut/Beatis_L3_1.fq.gz.1.gz

#1ba32f3d3a72be4877311163ce07ddc6 Beatis_L3_1.fq.gz.cut/Beatis_L3_1.fq.gz.2.gz

#26eaa5697b1c950aba2d83095f143f0c Beatis_L3_2.fq.gz.cut/Beatis_L3_2.fq.gz.1.gz

#f7fee1ecf2b928cf4504e6d48f417636 Beatis_L3_2.fq.gz.cut/Beatis_L3_2.fq.gz.2.gz

#Check per permissions, the following code changes the permissions of the file called et_trimmer.pl.

**chmod 777 est_trimmer.pl**

#Unzipping

#E.g. Rawdata files end in “gz” so they need to be unzipped:

#WTCHG_93433_274_2.fastq.gz

#WTCHG_93433_274_1.fastq.gz

#WTCHG_93434_274_1.fastq.gz

#WTCHG_93434_274_2.fastq.gz

#The following command will unzip everything ending in.gz in the background. The above file goes from fastq.gz file to just fastq file.

**gunzip *.gz &**

#For unzipping program files e.g. musket. If ends in tar.bz the command is:

**tar-xvjf**

#If ends in tar.gz the command is:

tar-xvzf

##################################################################

### 2.0. Concatenate

#I had two libraries with forward and reverse, put the two forwards into one file and the two backs in one file, just to make it simpler. The two 1’s together and the two 2’s.

#The following command tells Linux to concatenate the files called ‘WTCHG_93433_273_l.fastq’ and ‘WTCHG_93434_273_1.fastq’, and name the combined file iso_1.fastq, and do it all in the background (&). Note, you have to be in the directory that the files are in or tell Linux where to find them e.g. home/c1135170/Hannah/

**cat WTCHG_93433_273_1.fastq WTCHG_93434_273_1.fastq > iso_1.fastq &**

#Do the same with the other pair

**cat WTCHG_93433_273_2.fastq WTCHG_93434_273_2.fastq > iso_2.fastq &**

#E.g with *B. rhodani:*

**cat Beatis_L3_1.fq.gz.1 Beatis_L3_1.fq.gz.2 > B_1.fastq &**

**cat Beatis_L3_2.fq.gz.1 Beatis_L3_2.fq.gz.2 > B_2.fastq &**

##################################################################

### 3.0. Trimmomatic

#First need to download Trimmomatic, you can find here:

http://www.usadellab.org/cms/?page=trimmomatic

#Right click, ‘copy link ddress’ for the Binary.

**wget**

#right click to paste:

**http://www.usadellab.org/cms/uploads/supplementary/Trimmomatic/Trimmomatic-0.32.zip**

#It downloads to the folder you’re in and it’s called Trimmomatic-0.32.zip

**unzip Trimmomatic-0.32.zip**

#now just called Trimmomatic-0.32

#Now ready to run Trimmomatic

#Code means: Using Trimmomatic PE (paired end) which can be found here (pathway) do “‒phred33” to these two files (B_1.fastq & B_2.fastq) then rename them trimmomatic_B_1.fastq.gz (for paired) and trimmomatic_B_1_unpaired.fastq.gz for unpaired, and the same with the other pair. The nohup at the beginning is there so I can close the window and it will still carry on running.

**nohup java ‒classpath /home/c1135170/Hannah/app/Trimmomatic-0.32/trimmomatic-0.32.jar org.usadellab.trimmomatic.TrimmomaticPE ‒phred33 B_L.fastq B_2.fastq trimmomatic_B_1.fastq.gz trimmomatic_B_1_unpaired.fastq.gz trimmomatic_B_2.fastq.gz trimmomatic_B_2_unpaired.fastq.gz**

**ILLUMINACLIP:/home/c1135170/Hannah/app/Trimmomatic-0.32/adapters/TruSeq2-PE.fa:2:30:10 LEADINGS:3 TRAILING:3 SLIDINGWINDOW:4:18 MINLEN:35 &**

#E.g. Nohup.txt

#Read Pairs: 61528668 Both Surviving: 61132622 (99.36%) Forward Only Surviving: 96495 (0.16%) Reverse Only Surviving: 298538 (0.49%) Dropped: 1013 (0.00%)

##################################################################

### 4.0. Musket

#Have to download Musket the same way as Trimmomatic, you’ll find musket here:

http://musket.sourceforge.net/homepage.htm#latest

#Version used musket-1.1, right click, copy link address as with Trimmomatic:

**wget**

#right click to paste

**http://sourceforge.net/projects/musket/files/musket-1.1.tar.bz**

#When pasting link may have “/download” on the end of the link, delete this before pressing enter.

#Unzip file (see section 1.0)

#To install, go to the Musket folder and press ‘make’:

**cd ‥/app/cd musket-1.1/is**

**make**

#Then you can remove the original zipped file

**rm musket-1.1.tar.bz**

#2.0. Run Musket using file outputs from Trimmomatic:

**nohup /home/c1135170/Hannah/app/musket-1.1/musket ‐k 21 2192141955 ‐p 32 ‐omulti correctedinorder trimmomatic_B_1.fastq.gz trimmomatic_B_2.fastq.gz trimmomatic_B_1_unpaired.fastq.gz trimmomatic_B_2_unpaired.fastq.gz 1>out.txt 2>error.txt**

#They are named corrected.0, corrected.1, corrected.2, and corrected.3, in order of how the files were listed in the command above. Rename the files:

**mv corrected.0 musket_B_1.fastq**

**mv corrected.1 musket_B_2.fastq**

**mv corrected.2 musket_B_1_unpaired.fastq**

**mv corrected.3 musket_B_2_unpaired.fastq**

##################################################################

### 5.0. FLASH (used for *O. asellus only*)

#To find Flash: http://ccb.jhu.edu/software/FLASH/

#To download

**wget** http://sourceforge.net/proiects/flashpage/files/FLASH-1.2.9.tar.gz

#To unzip

**tar ‐zxvf FLASH-1.2.9.tar.gz**

\# Command asks flash to merge paired ends <musket_woo_1> < musket_woo_1> [-m minOverlap - varied] [-M maxOverlap-100] [-x mismatchRatio-varied] [-p phredOffset] [-0 prefixOfOutputFiles] [-d pathToDirectoryForOutputFiles] [-f averageFragment Length-300] [-s standardDeviationOfFragments-varied] [-r averageReadLength-150].

#Several different compbinations tried to

**‥/app/FLASH-1.2.9/flash musket_woo_1.fastq musket_woo_2.fastq ‐m 15 ‐M 100 ‐x 0.1 ‐p ‐o merged ‐d ‐f 300 ‒s 50 ‐r 150 1>flash.out 2>flash.err &**

#10% retained and matched

**‥/app/FLASH-1.2.9/flash musket_woo_1.fastq musket_woo_2.fastq ‐m 10 ‐M 100 ‐x 0.1 ‐o merged2 ‐d ‐f 300 ‒s 40 ‐r 150 1>flash2.out 2>flash2.err &**

#15% retained and matched

**‥/app/FLASH-1.2.9/flash musket_woo_1.fastq musket_woo_2.fastq ‐m 25 ‐M 100 ‐x 0.1 ‐o merged3 ‐d ‐f 300 ‒s 40 ‐r 150 1>flash3.out 2>flash3.err &**

#9.5% retained and matched

**‥/app/FLASH-1.2.9/flash musket_woo_1.fastq musket_woo_2.fastq ‐m 20 ‐M 100 ‐x 1 ‐o merged4 ‐d ‐f 300 ‐s 40 ‐r 150 1>flash4.out 2>flash4.err &**

#100% retained and matched

**‥/app/FLASH-1.2.9/flash musket_woo_1.fastq musket_woo_2.fastq ‐m 20 ‐M 100 ‐o merged4 ‐d ‐f 300 ‐s 40 ‐r 150 1>flash4.out 2>flash5.err &**

#mismatchRatio: default 0.25.12.78% retained and matched

#Use merged extendedfrags to feed straight into MISA

#To continue must convert fastq file to fasta file, and removes spaces at the same time, using:

**awk'BEGIN{a=0}{if(a==l){print;a=0}}/⌃@/{print;a=l}' myFastqFile | sed 's/⌃@/>/' > myfastafile**

##################################################################

### 6.0. SOAPdenovo2

#Download soapdenovo, (same way as Trimmomatic and Musket), find here:

http://sourceforge.net/proiects/soapdenovo2/files/SOAPdenovo2/

#To download:

**wget** http://sourceforge.net/proiects/soapdenovo2/files/latest/download?source=files

#Unzip

**tar ‐xvvf SOAPdenovo2-src-r240-4.tar**

#Compile by navigating to the folder that the 'makefile' is in and type:

**Make**

#First make config file to use with soap de novo, red needs to change depending on the data, especially the pathways to the musket output files so soap de novo knows where the files are. In this example the config file was named ‘iso_config.text’.

max_rd_1en=**150**

[LIB]

#average insert size

avg_ins=**300**

#if sequence needs to be reversed

reverse_seq=0

#in which part(s) the reads are used

asm_flags=3

#in which order the reads are used while scaffolding

rank=1

#a pair of fastq file, read 1 file should always be followed by read 2 file

q1=/home/c1135170/**Hannah/Isoplera/ musket_iso_1.fastq**

q2=/home/c1135170/**Hannah/Isoplera/musket_iso_2.fastq**

q=/home/c1135170/**Hannah/Isoplera/musket_iso_1_unpaired.fastq**

q=/home/c1135170/**Hannah/Isoplera/musket_iso_2_unpaired.fastq**

#For kmers less than 63, use the following command, be in the same folder as the config file:

**nohup ‥/app/soapdenovo/SOAPdenovo2-src-r240)/SOAPdenovo-63mer all ‒s iso_config.txt - K 55 ‐R ‐o iso_kmer55 1>iso_kmer55.log 2>iso_kmer55.err**

\# Command explained: **nohup ‥/app/soapdenovo/SOAPdenovo2-src-r240)/SOAPdenovo-63mer** (telling it where to find soapdenovo) **all ‒s iso_config.text** (name of the config file we made) **‒K 55** (kmer size 61)**–R ‒o iso_kmer55 1>iso_kmer55.log 2>iso_kmer55.err** (names of the output files)

#For Kmers above 63:

**Nohup /home/c1 13517C)/Hannah/app/SOAPdenovo2-bin-LINUX-generic-r24C)/SOAPdenovo-127mer all ‐s iso_config.txt ‐K 71 ‐p 10 ‐R ‐o iso_kmer71 1>kmer71.log 2>kmer71.err**

